# A crescendo of competent coding (c3) contains the Standard Genetic Code

**DOI:** 10.1101/2022.05.22.492986

**Authors:** Michael Yarus

**Affiliations:** Department of Molecular, Cellular and Developmental Biology, University of Colorado Boulder, Boulder, Colorado 80309-0347

## Abstract

The Standard Genetic Code (SGC) can arise by fusion of partial codes evolved in different individuals, perhaps for differing prior tasks. Such code fragments can be unified into an SGC after later evolution of accurate third-position Crick wobble. Late wobble advent fills in the coding table, leaving only later development of final translational initiation and termination in separate domains of life. This code fusion mechanism is computationally implemented here. C3 fusion before late Crick wobble (c3-lCw) is tested for its ability to evolve the SGC. Compared with the previously-studied evolution of isolated coding tables, or with increasing numbers of similar, but non-fusing codes, code fusions reach the SGC sooner, work in a smaller population, and present more accurate and more complete codes more frequently. Notably, a crescendo of SGC-like codes is exposed to selection for an extended period. c3-lCw also effectively suppresses varied disordered assignments, unifying the coding table. Such codes approach the SGC, making its selection seem likely. Given unexceptional conditions, ≈ 1 of 22 c3-lCw environments evolves codes with ≥ 20 assignments and ≤ 3 differences from the SGC, including some with assignments identical to the SGC.

## Introduction

The Standard Genetic Code associates 22 functions (20 amino acids plus initiation and termination) with the 64 possible ordered RNA triplets in a way reproduced with appreciable accuracy throughout Earth’s biota. This implies that the SGC preexisted in ancestors of all modern Earth creatures. Thus, the SGC’s derivation offers information about early biology before the common ancestor of modern organisms, and during divergence into present (Zhou et al. 2018) domains of life.

Here, such information is sought by quantitative modeling of SGC emergence, using, arguably, general assumptions (Yarus 2021b). It is assumed only that codon assignment, capture and decay (and added here: new coding tables and code fusion) occur at characteristic rates. SGC existence is attributable to the joint effects of those rates within the 64 triplet space of a coding table.

In order to embody events whose complexity may be great, but unknown at the start, a computable model is used (Yarus 2021b; Fig. 7). This envisions SGC evolution as a set of shorter intervals, called passages, during which one event only (a codon assignment, for example, or possibly no event) occurs. Encoding events differ in probability during a passage. The virtue of this formulation is that it allows explicit programming of hypotheses about code evolution, even for histories of great complexity. Implied coding tables are readily computed. Moreover, using different probabilities during an interval is equivalent to assignment of different rate constants (Yarus 2021b), and timing of events may therefore be compared. For example, when the real-world time to assign a codon is known, these calculations convert to times on an early Earth (Yarus 2021d).

These inquiries take their most recent form (Yarus 2021c) as the idea that the code was composed by fusing independently-developed partial codes, perhaps fusing primitive compartments that had developed differing coding competencies. Code fusion is common in Biology, having been observed many times, for example, between mitochondrial and nuclear codes (Duchêne et al. 2009).

Creation of the SGC by fusion of separate partial codes was thought (Yarus 2021c) to have specific advantages; for example, realizing the SGC within a smaller code population. Below, hypothetical advantages of code fusion during SGC construction are tested.

## The model

### Individual coding tables

Developing primordial codes (Yarus 2021b) may assign unassigned triplets (with probability Pinit), using either SGC-like assignments or random assignments (probability Prand), or capture unassigned triplets related by single mutations from their existing assignments (Pmut). Such assignments, however, can decay and be lost (Pdecay). Probabilities have the same relative values for passages here as in prior studies, so present codes resemble those earlier ones. However, probabilities have also been reduced in proportion so that two events in one interval are less probable. This makes the present model somewhat more accurate, though more passages elapse during code evolution.

### Fusion of codes

In addition, new events (Fig. 7) occur in this work: with constant probability/passage, new coding tables appear (Ptab). New codes begin with a single arbitrary assignment (Pinit), then evolve using the same rules as the initiating coding table. Newly originated codes accumulate; once these exist, they may fuse with other codes (Pfus) with a probability that increases with the number of possible partners. A fused compartment can gain assignments from both fusees, or it can be unchanged, if both happen to use overlapping prior encodings. However, fusion can also be disastrous, if fusing codes conflict. A simple rule is used: if fused codes give a triplet more than one meaning, this will be damaging, and both participants are lost.

### The evolutionary goal

The assumption is: there was a functional advantage to SGC coding, and codes more like it increasingly possess that advantage. To avoid unnecessary hypotheses, superiority of the SGC is unspecified. Instead, code selection is more probable as the distance to the SGC decreases. This is implemented by seeking codes that are sufficiently complete: they encode >= 20 of the 22 possible functions (recognizing the late development of definitive initiation and termination (Burroughs and Aravind 2019)). In addition, codes must be accurate: they vary from the SGC by the fewest misassignments, abbreviated “misx”, where x is the number of differences from the SGC. Codes closest to SGC completeness and SGC assignments (called SGC-like codes) are most likely to have been selected, whatever (unknown) selection may have applied.

### Fusion must be a major evolutionary event

Coding evolution is altered by fusion only if it occurs significantly. This frequency requirement is both elementary and profound; we discuss fusion rates first.

Code fusion requires two successive events. Firstly, new coding tables arise to create a population of codes in the initial code’s environment. This happens at a fixed probability per passage (Ptab); a passage being mean time for completion of one evolutionary step for a coding table.

In the second step, tables, once multiple ones exist, may fuse their codes at random, with probability Pfus per passage, Pfus*(others). (others) is the number of codes existing alongside each fusion candidate; thus (others) = (total codes - 1). Fusions become increasingly probable with time.

Thus, time is measured in passages, which simultaneously host evolution in all existing codes (Methods). Either a new assignment, capture of a sequence-related codon, assignment decay, or a code fusion may occur within a passage, perhaps accompanied by creation of a new coding compartment that begins with one assigned codon, and evolves alongside the initial code.

As an environment’s code population increases, it acquires more complex coding, and the program records these events in any detail desired. Every change and every intermediate code can be recorded and delivered to the experimenter. But because only a small part of total change is usually of interest, only partial data are routinely reported. The first is a summary of the important properties of every code in an environment at every passage. This allows study of average events for all codes. The second kind of report tracks only most complete codes (e.g., most functions assigned). This pursues advanced coding, useful if progress to SGC-like codes is being studied.

### A majority of fusions

Fig. 1 plots the fraction of codes with >= 20 assigned functions that have benefitted from a fusion contribution, versus the probability of new tables (Ptab in Fig. 1A, Pfus is fixed and favorable), or versus the probability of fusion (Pfus in Fig. 1B, Ptab is fixed and favorable). More tables and more likely fusion, as expected, increase the fraction of SGC-like (>=20 function) codes that acquire fused assignments. Because this ms concerns change produced by code fusions, Ptab = 0.08 and Pfus = 0.002 (Fig. 1, rightward, maximal fusion) are used below. A requirement for elevated fusion parallels other studies (Aggarwal et al. 2016), and provides a simple rationale: fusion must be a significant route to final codes.

**Figure 1A.**
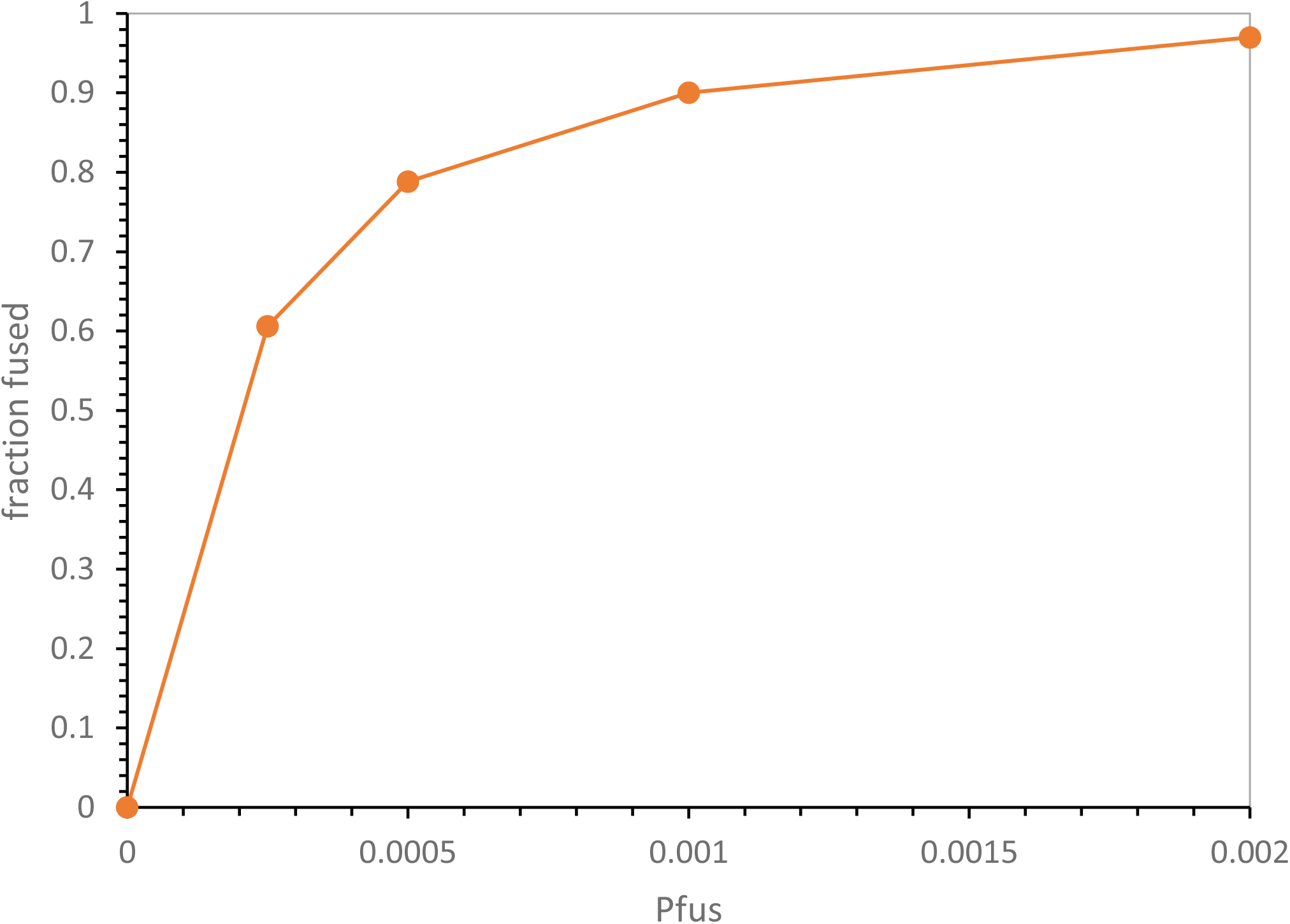
Complete fused codes vs probability of fusion. Evolution was stopped when the first code with >= 20 functions appeared in an environment. The fraction, among 500 environments, of such >= 20 function codes that acquired assignments from fusion is plotted versus the probability of fusion during a passage, Pfus. Pmut = 0.00975, Pdecay = 0.00975, Pinit = 0.150, Prand = 0.050, Ptab = 0.08.

**Figure 1B.**
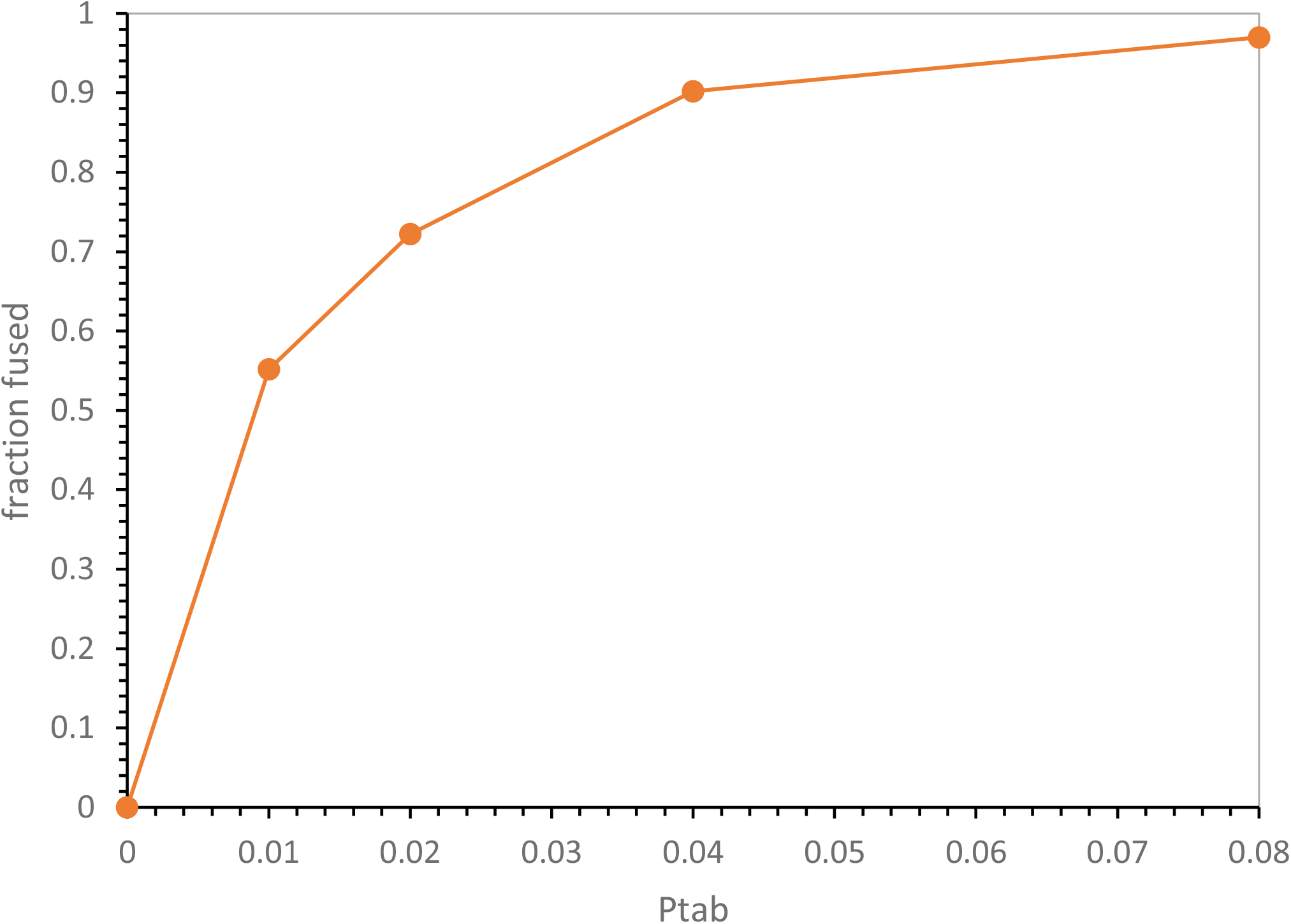
Complete fused codes vs probability of coding table initiation. Evolution was stopped when the first code with >= 20 functions appeared in an environment. The fraction, among 500 environments, of such >= 20 function codes that acquired assignments from fusion is plotted versus the probability that a new cooding table is initiated during a passage, Ptab. Pmut = 0.00975, Pdecay = 0.00975, Pinit = 0.150, Prand = 0.050, Pfus = 0.002.

### Altered population history

Fig. 2 contains averages for every coding table in 1000 code populations carried to 750 passages. Beginning with a single initial code, this ranges from 5412 potential codes at 60 passages to 60790 potential codes at 750 passages. Thus, accurate mean values for code kinetics are available. In Fig. 2, all coding tables are partitioned into 4 classes - they have had no fusion, have been lost in fusion, have been annihilated by incompatible fusion, or have received successful fusion (Fig. 7).

**Figure 2.**
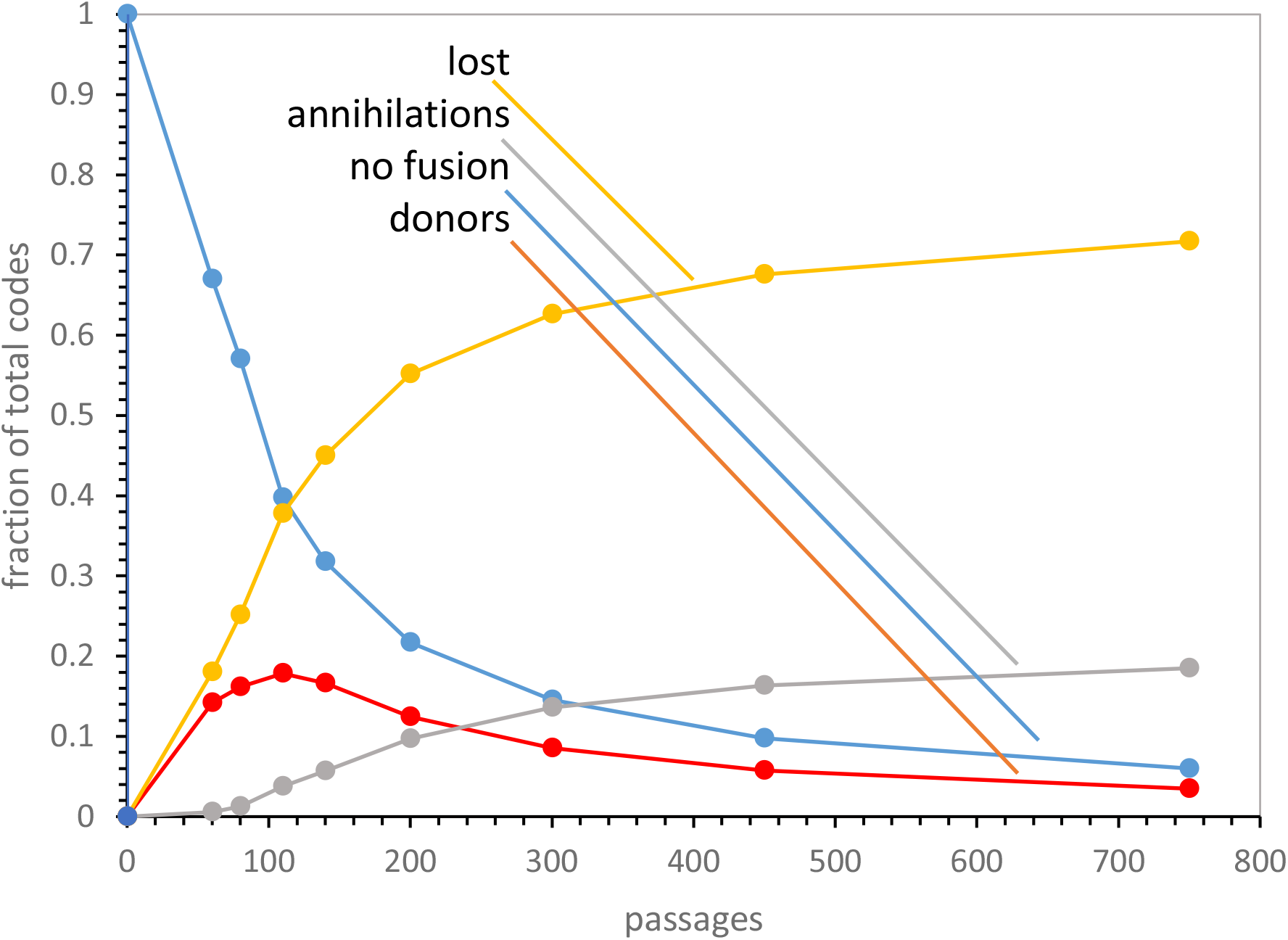
Code fate vs time. Mean fractions of 1000 total codes are plotted versus passages (time). Kinetics for several fates are shown: donors – codes that successfully fused / annihilations – codes lost in incompatible fusions / no fusion – unfused codes / lost – donors lost in fusion. Pmut = 0.00975, Pdecay = 0.00975, Pinit = 0.150, Prand = 0.050, Pfus = 0.00200, Ptab = 0.08.

These times allow unfused tables to become infrequent (only about 6% are unfused at 750 passages). The predominant fate of coding tables is loss in fusion, and this is true from early times, just before 140 passages. Codes are lost because they fuse into others, and a significant minority were annihilated by trying to fuse to codes with incompatible assignments. Putting the same data in other words, only 9.4% of once-existent codes still exist at 750 passages (those with no fusion or successful fusions). Successful fusions themselves have a peak around 110 passages, after which they are also lost in later destructive events. Fig. 7, which sketches kinetics for a simplified environment, may help conceptualize fusion losses. We will return to the early successful fusion maximum (Fig. 2), and to its later decline, below.

### Superior codes follow fusion

We now examine later codes in Fig. 2; this minority of fusion survivors includes codes that closely approach the SGC. Fig. 3 shows the properties of the most complete codes from 10000 code populations evolved throughout the interesting era of Fig. 2, from 150 to 750 passages, when fusions emerge, then decisively shape, an environment’s codes.

**Figure 3A.**
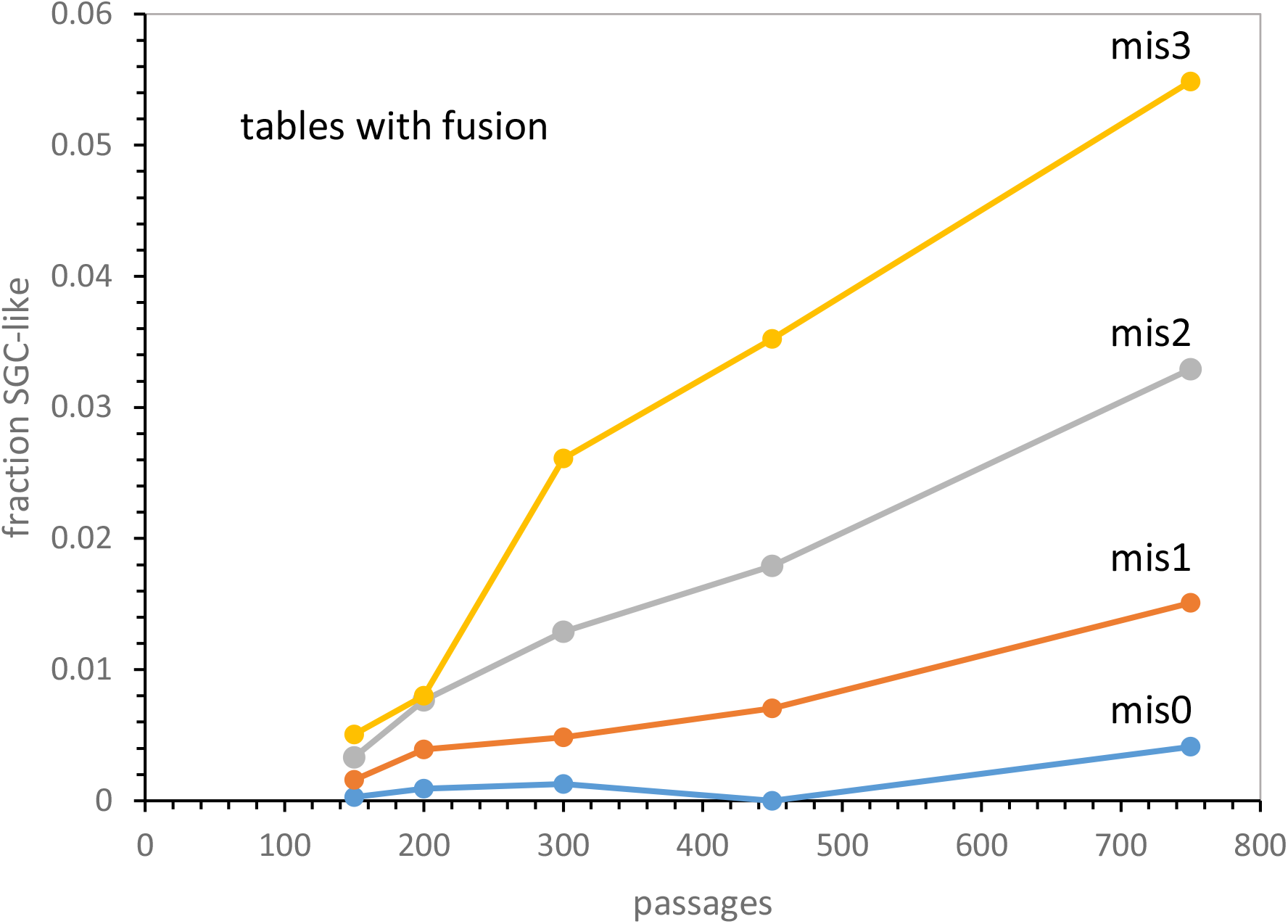
SGC-like codes vs time: tables and fusion. The fraction of almost complete codes (>= 20 functions) with cited levels of misassignment (relative to the SGC) is plotted for 10000 environments that have both new code initiation and fusion. mis0 = no misassignments / mis1 = 1 misassignment, and so on. 10000 environments evolved to the times/passages shown, and codes with >= 20 assigned functions were characterized. Pmut = 0.00975, Pdecay = 0.00975, Pinit = 0.150, Prand = 0.050, Pfus = 0.002, Ptab = 0.08.

### Competence

Fig. 3A depicts abundance of SGC-like codes. These codes have either experienced no fusion, or a complementary fusion that adds assignments. They have assigned codons to >= 20 of the standard functions, and so are almost complete. Moreover, plotted compartments encode functions very similarly to the SGC, having assignments completely overlapping (blue line), a single differing assignment (red), two differences (gray) or 3 differently-assigned triplets (yellow).

### The crescendo

The fraction of competent codes tends to rise from an origin just after the appearance of population-wide fusions (Fig. 2) to the end of calculation (Fig. 3A; the crescendo). In addition, these triply-unusual codes, with completeness, fusion contributions and accuracy that are all exceptional, are quite frequent, seemingly well within the reach of a search for SGC-like translation. For example, the completely SGC-like class (mis0) are detected in 10000 environments early, at 150 passages, and are 1/250 among the best codes at 750 passages. Even supposing a demanding coding selection, requiring precise SGC mimicry, such codes appear relatively early, and require selecting a superior translational system in ≈ 1/250 environments. This seems achievable.

Further, if selection for superior translation extends to all codes similar to the SGC, >= 20 function, mis0 to mis3 codes exist in 1/100 environments at 150 passages, and more frequently than 1/10 environments at 750 passages. Late selection for superior encodings would not seek far for SGC-like results.

### The crescendo evolves

Existing codes (Fig. 3A, 3B, 3C) quickly acquire additional assignments. Less quickly, new codes arise and existing codes fuse. Less frequently yet, new codons are captured for existing assignments and assignments decay. Thus, a large flux of change is absorbed by evolving codes. The implication is that the competent code population is constantly changing on a timescale comparable to its initial appearance. The coding crescendo’s tables are constantly evolving, with new codes replacing the previously competent: compare successful fusions lost to later events (Fig. 2). Thus the crescendo offers a changing face to selection, as well as increasingly frequent competent codes. Selection can occur when a particularly effective SGC-like code appears in the crescendo’s jumble.

**Figure 3B.**
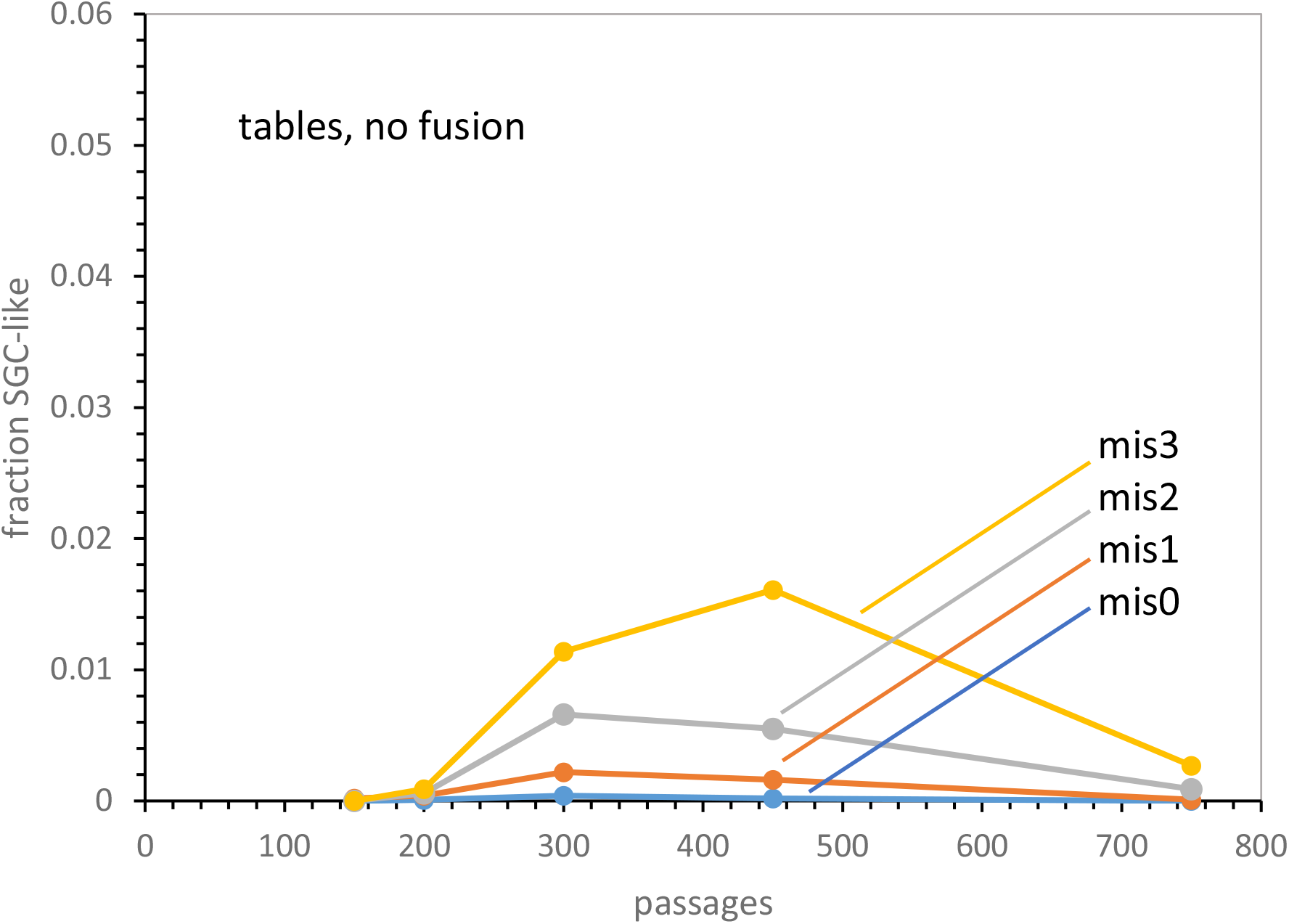
SGC-like codes vs time: tables, no fusion. The fraction of almost complete codes with cited levels of misassignment (relative to the SGC) is plotted for 10000 environments that have new code initiation, but no fusion. mis0 = no misassignments / mis1 = 1 misassignment, and so on. 10000 environments evolved to the times/passages shown, and codes with >= 20 assigned functions were characterized. Pmut = 0.00975, Pdecay = 0.00975, Pinit = 0.150, Prand = 0.050, Pfus = 0.000, Ptab = 0.08.

**Figure 3C.**
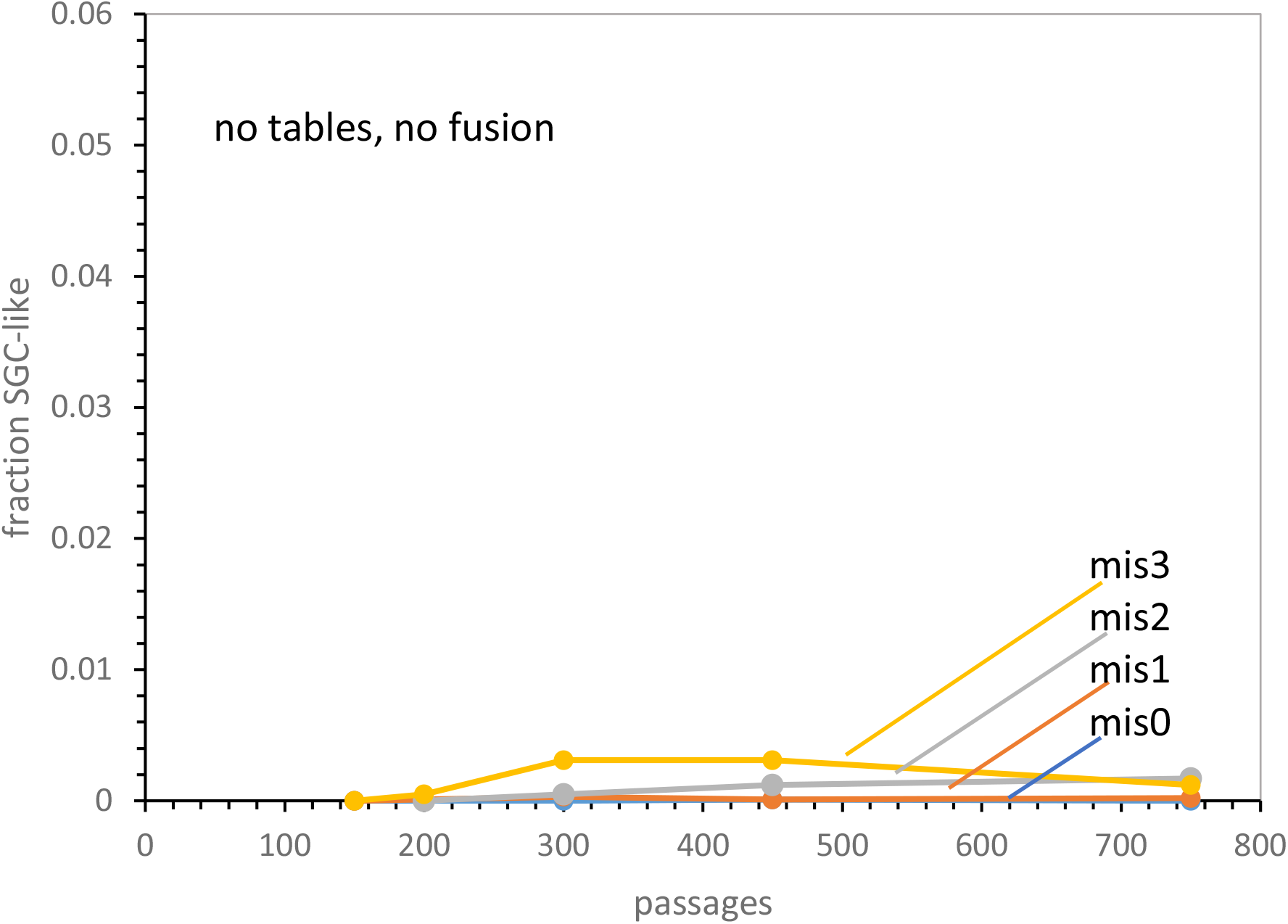
SGC-like codes vs time: no new tables, no fusion. The fraction of almost complete codes with cited levels of misassignment (relative to the SGC) is plotted for 10000 environments that have no new code initiation, and no fusion. mis0 = no misassignments / mis1 = 1 misassignment, and so on. 10000 environments with single tables evolved to the times/passages shown, and codes with >= 20 assigned functions were characterized. Pmut = 0.00975, Pdecay = 0.00975, Pinit = 0.150, Prand = 0.050, Pfus = 0.000, Ptab = 0.00.

### The crescendo and its competence come from fusion I

The crescendo of competence is produced by code fusion.

Fig. 3B shows code evolution similar to Fig. 3A, but without fusion (Pfus = 0). As in other Fig. 3 panels, fractions of the most complete codes from 10000 environments (Fig. 12) are plotted versus passages (time). Having multiple codes itself facilitates the evolution of more complex coding. Thus environments with parallel coding tables as in Fig. 3A, but with no fusions between them (Fig. 3B), present a useful comparison. Fig. 3B is plotted with the same y-axis to facilitate comparison with Fig. 3A: no crescendo exists. In fact, for multiple tables without fusion, all levels of completeness with assignment accuracy arise later, achieve lower frequencies among most complete codes, and do not persist. Competence ultimately declines instead of increasing (Fig. 3A).

### The crescendo and its competence come from fusion II

Multiple coding tables facilitate evolution of the SGC without fusing. Fig. 3C completes controls for Fig. 3A; it describes a similar set of code evolutions, but lacking both multiple tables and fusion (Ptab = 0, Pfus = 0). This resembles the system previously analyzed (Yarus 2021b), where code evolution takes place in a single initial coding table in each environment, each evolving until it resembles the SGC. However, Fig. 3C is useful here because its individual codes are the same as those of Fig. 3A and 3B. Fig. 3C also has the same time scale, frequency scale and colors as Fig. 3A and 3B. Thus it is clear that single codes without fusion gain SGC resemblance later than in Fig. 3A, restrict such competence to lower levels even than for multiple tables in Fig. 3B, and again show no crescendo (Fig. 3A). In fact, SGC-like coding is everywhere lower than for multiple tables without fusion (Fig. 3B). For example, complete resemblance to the SGC (mis0) is not detected among 10000 environments until 450 passages and then at too low a frequency (≈ 10^−4^) to be deciphered on Fig. 3C’s ordinate.

### Origin of competence

It is clear why environments that fuse code compartments, cells or partial coding tables are superior. Fig. 4A compares mean misassignments for the most complete codes (having assigned >= 20 functions) from 10000 environments. Multiple codes with fusion (red, Fig. 4A) are similar to multiple codes without fusion (green, Fig. 4A) and to codes without multiple tables and fusion (blue, Fig. 4A) until fusion becomes predominant in code evolution (see **Altered population history**, above). After this time (≈ 150 passages, Fig. 4A) errors during different evolutionary modes diverge greatly. Strikingly, other modes almost double mean misassignment in codes with fusion.

**Figure 4A.**
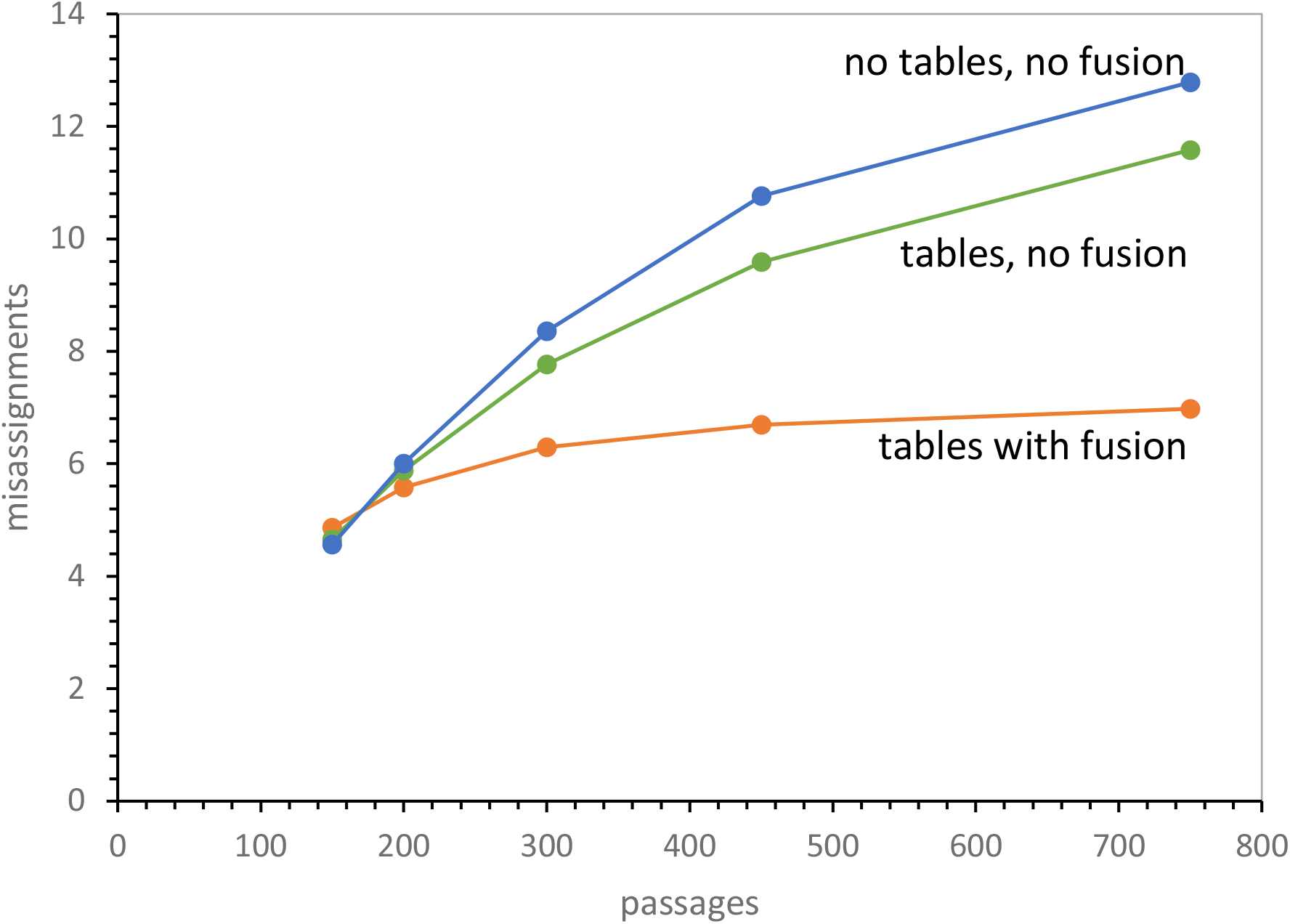
Mean misassignments vs time: tables and fusion, tables no fusion, no tables no fusion. 10000 environments were evolved to the times shown, and misassignments relative to the SGC were counted among most complete codes in each environment. Probabilities are the same as in Fig. 3 for the three kinds of evolution.

There are two sources of misassignment in present code environments. The more straightforward is that random assignment is allowed by a variable probability of random association between functions and triplets (Prand). Because these assignments are accurately randomized, they are unlikely to be the same as for the SGC.

### Fate of random assignments

Fig. 4B shows the mean number of randomized assignments in the most complete codes from 10000 coding environments (Prand = 0.05) versus time. The pattern strikingly reproduces overall accuracy in Fig. 4A. That is, codes derived by fusing multiple tables (blue, Fig. 4B) have made 2-3 fold fewer random assignments than without fusion (red, Fig. 4B) or without both multiple tables and fusion (gray, Fig. 4B).

**Figure 4B.**
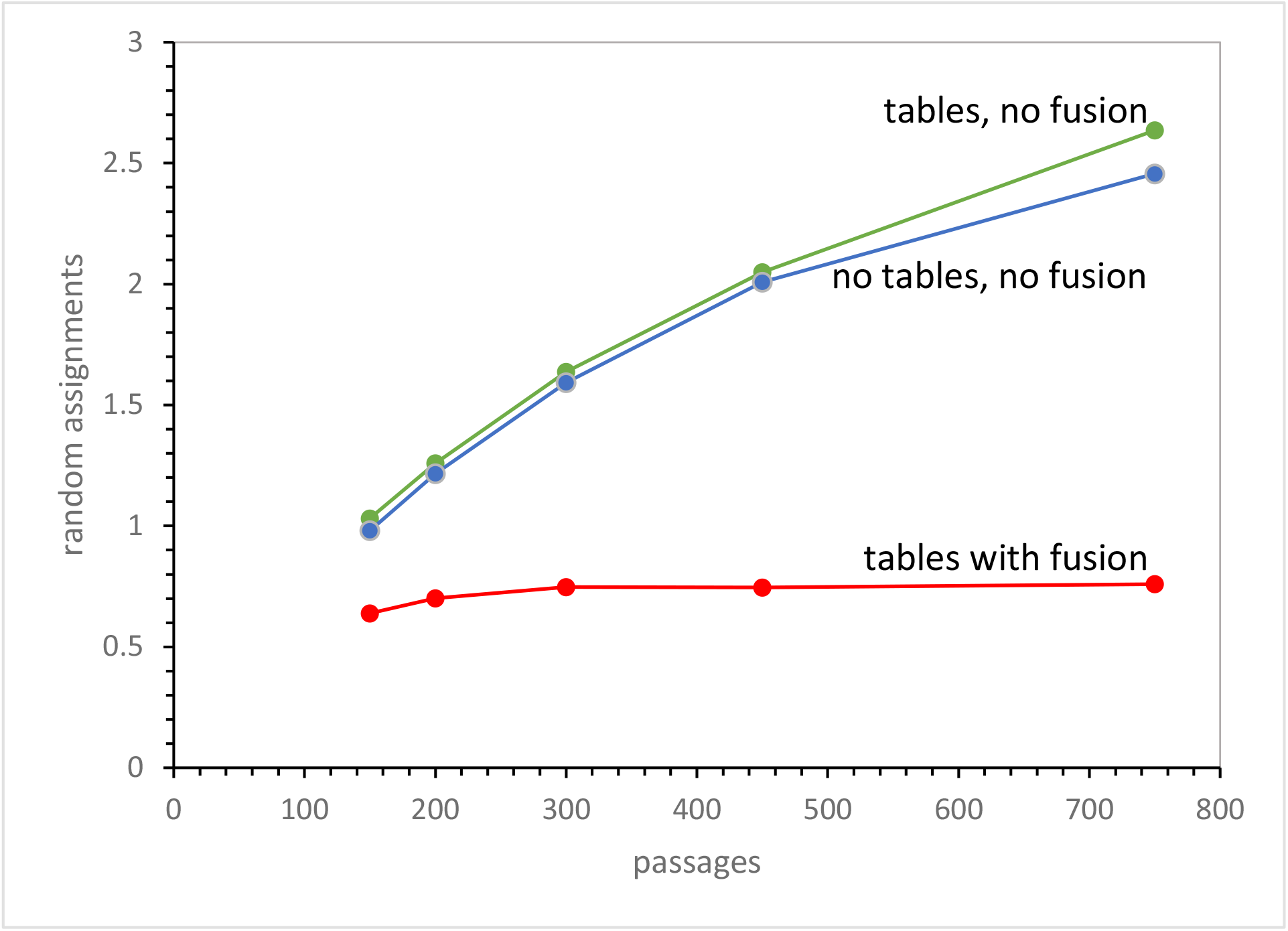
Randomly assigned codons vs time: tables and fusion, tables no fusion, no tables no fusion. 10000 environments were evolved to the times shown, and randomly assigned codon triplets were counted among most complete codes in each environment. Probabilities are the same as in Fig. 3 for the three kinds of evolution.

### Fate of captured triplets

The second source of misassignment is that related triplets (one mutation away from an assigned triplet) can be captured for an existing related function. The new assignment can be to a chemically related amino acid (having similar polar requirement (Mathew and Luthey-Schulten 2008; Woese 1965)), or even the same as the previously assigned function (Yarus 2021b). Chemically related amino acids are sometimes, but not always, assigned to mutationally related triplets in the SGC, and there are several choices for the ‘chemically related’ one (see Methods). So, capture also frequently yields encoding unlike the SGC.

Fig. 4C shows that codes using fusion more strongly discriminate against captures of mutationally related triplets for related functions. Again, the pattern follows that in Fig. 4A: before prevalent fusion, the three modes of code evolution are similar. Afterward, they progressively diverge under fusion; at 750 passages fused codes (blue, Fig. 4C) utilize 4-5 fold fewer error-prone captures than do unfused multiple codes (red, Fig. 4C) or single codes (gray, Fig. 4C).

**Figure 4C.**
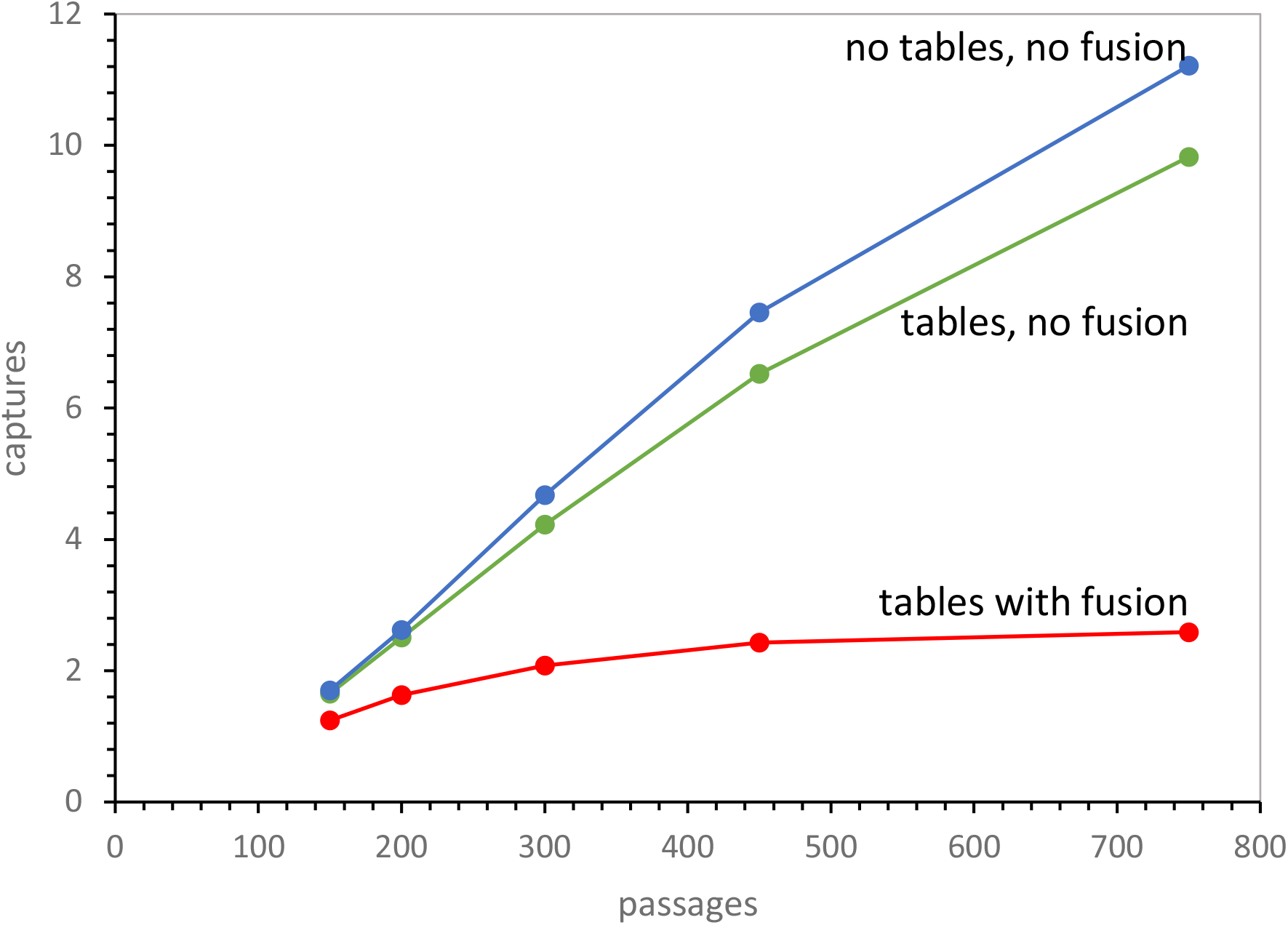
Capture of mutationally related codons vs time: tables and fusion, tables no fusion, no tables no fusion. 10000 environments were evolved to the times shown, and capture of triplets one mutation distant from assigned codons were counted among most completely assigned codes in each environment. Probabilities are the same as in Fig. 3 for the three kinds of evolution.

### Codes with misassignments are rejected

How do fused codes become superior? Fusion tests codes against each other because unlike codes are incompatible. Fusions between like codes are more likely to succeed; fusions between unlike codes are more likely to be lost because of the toxic effects of codons with multiple meanings. Thus when highly complete fused codes are characterized above (Fig. 3, 4), they are intrinsically less heterogeneous than the partial codes from which they have been derived. As a fusing environment progresses, with fusion more and more probable (Fig. 2), heterogeneity due to random assignment (Fig. 4B) and capture of related codons (Fig. 4C) is suppressed among the fused (Fig. 4A). Therefore, the same number of unfused codes in one environment (Fig. 3B) or especially single codes without potential fusion (Fig. 3C) cannot compete with the accuracy of fusing tables evolving together (Fig. 3A).

### Acceptable randomness

The SGC is strikingly ordered, that is, non-random (Woese 1965). An important question for any coding scheme is therefore: how much random assignment can be tolerated? Increased competence via fused nascent codes specifically raises the possibility that fusion increases the latitude allowed to early coding. Fig. 5 thus presents data for random assignment from none (Prand = 0) to about 2.2 random assignments/code on average (Prand = 0.1).

**Figure 5.**
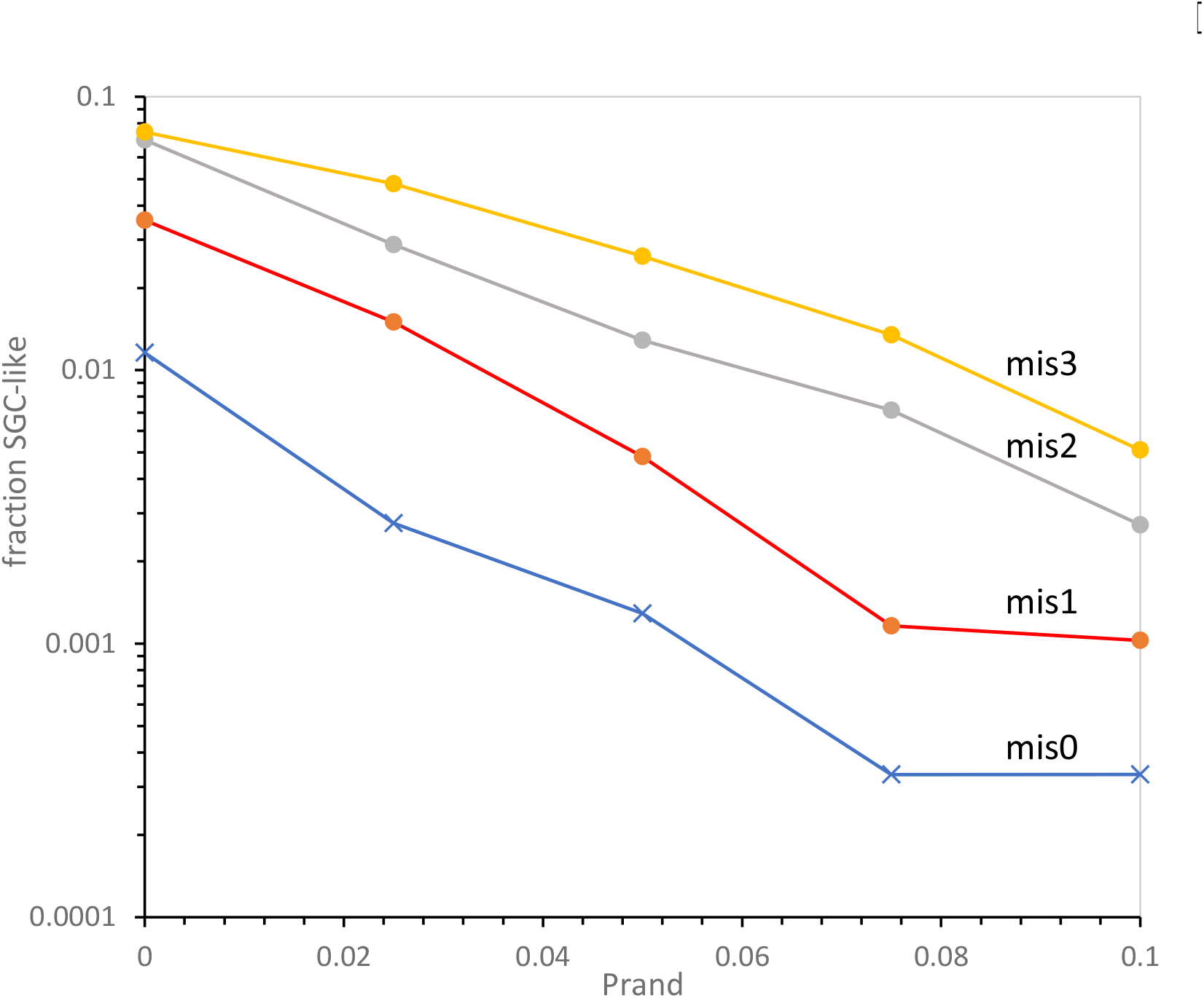
Fraction SGC-like codes vs probability of random assignments. 10000 (or 20000 for greatest Prand) environments were evolved to 300 passages. Among substantially complete codes (>= 20 assigned functions), fractions with different levels of misassignment were counted. mis0 = no misassignments relative to the SGC / mis1 = 1 misassignment, and so on. Pmut = 0.00975, Pdecay = 0.00975, Pinit = 0.150, Pfus = 0.00200, Ptab1 = 0.08.

Data in Fig. 5 are for tens of thousands of environments at 300 passages, a time when all modes are evolving SGC-like codes (Fig. 3). The frequencies of near-complete, accurately assigned codes are plotted logarithmically versus the linear probability of random assignment (Prand). Log frequencies of such SGC-like codes tend to decrease linearly with Prand (see also (Yarus 2021c)), with decrease somewhat more rapid as the rigor of requirement increases. Thus, inerrant codes (mis0) decrease somewhat more than those with three misassignments (mis3). But even if Prand = 0, coding is not perfect, because other sources of error remain, like capture of mutationally-related triplets for similar assignments (Fig. 4). Frequencies for good coding shown are higher than we have previously observed. These origin hypotheses still have substantial access to the SGC even if they randomly assign triplet functions in roughly 1 of 10 cases. Thus, the prior rule-of-thumb (Yarus 2021b) need not change for fused code evolution: 1 or 2 functions can have been assigned for different reasons, or for no specific reason.

## Discussion

### The Standard Genetic Code

Because all modern Terran organisms possess the SGC or a close relative, the most economical hypothesis is that the SGC already existed in the last common ancestor. The SGC’s origin has therefore attracted an immense literature that defies summary, especially within a single manuscript. However, below is a brief synopsis in order to put present findings in context.

### Optimization of the SGC

A large quantitative literature exists on optimization of the SGC, much of it initiated by the finding that code structure appears to reduce destructive effects of likely mutational or translational errors (Freeland and Hurst 1998). However, the SGC is only partially optimized (Novozhilov et al. 2007), standard optimization routes require many steps (Massey 2010), apparent optimization to errors readily occurs as a by-product of other goals (Massey 2008; Błażej et al. 2018) and full optimization is not physically plausible (Yarus 2021b). Thus, no purposeful code optimization exists in c3-lCw, other than that intrinsic to code fusion.

### Late Crick wobble (lCw)

Accurate third-codon-position position wobble (Crick 1966) is unlikely to be a primordial form of genetic coding. tRNA-rRNA interaction on the modern ribosome is extensive, spanning both tRNA molecules and sites in both large rRNAs (Moazed and Noller 1986, 1989). Some of these contacts appose rRNA nucleotides with codon-anticodon triplets to check their conformations (Ogle et al. 2001). Such checks determine whether the first two base pairs are Watson-Crick (Demeshkina et al. 2012), as well as whether wobble positions lie within the multiple conformations allowed for normal, tautomeric and charged wobble pairs (Westhof et al. 2019). Such sophisticated 3-dimensional error checking is unlikely for primordial encoding, but could evolve later in coding history. Thus, here coding is assumed to involve normal base pairing until a late time when wobble becomes possible, probably in a ribonucleopeptide proto-ribosome. Given that Crick wobble readily fills the coding table (Yarus 2021b), simplified Crick wobble is here assumed to be adopted quickly throughout a nascent SGC once it becomes possible. Others have also treated wobble as a significant late coding development (Lei and Burton 2021).

Late Crick wobble is implemented in explanatory diagrams (Fig. 6, 7), but actually plays little part in discussion of code capabilities because completeness (all functions encoded) is the usual criterion for code progress, and completeness is unaffected by Crick wobble, which only extends existing assignments.

**Figure 6.**
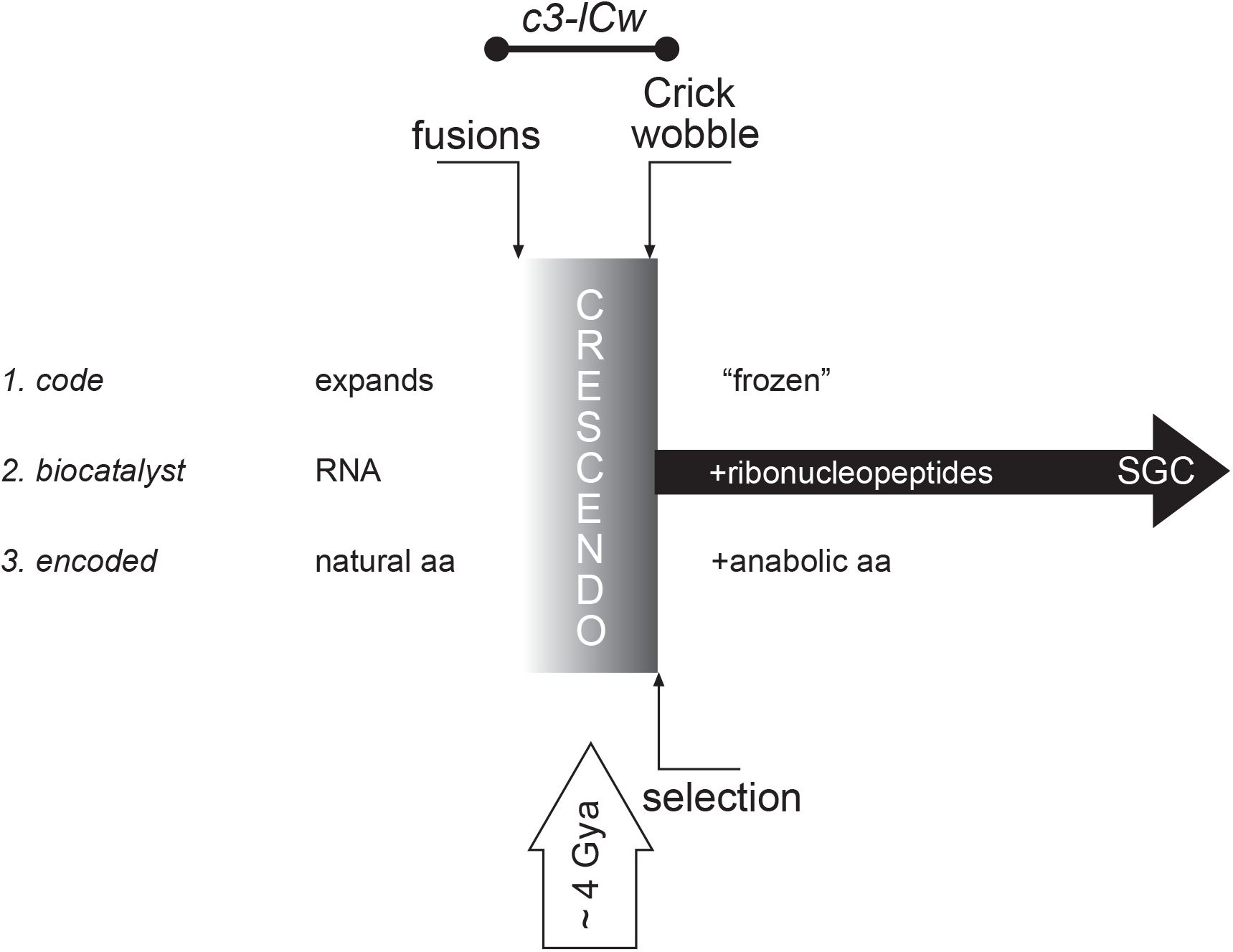
A coding crescendo separates two early code eras. Three numbered definitions for divided early code evolution (see text) are interpreted as outcomes of c3-lCw. Initiation of code fusion (“fusions”), evolution of wobble (“Crick wobble”) and selection of the SGC from the crescendo (“selection”) are marked relative to the crescendo (Fig. 3A). Late wobble evolves just before SGC selection, because wobble appears in the SGC. The last common ancestor lies off to the right of Fig. 6. The c3-lCw’s approximate time before the present (bottom, upward arrow) reflects that of the most ancient fossil biota (see text).

**Figure 7.**
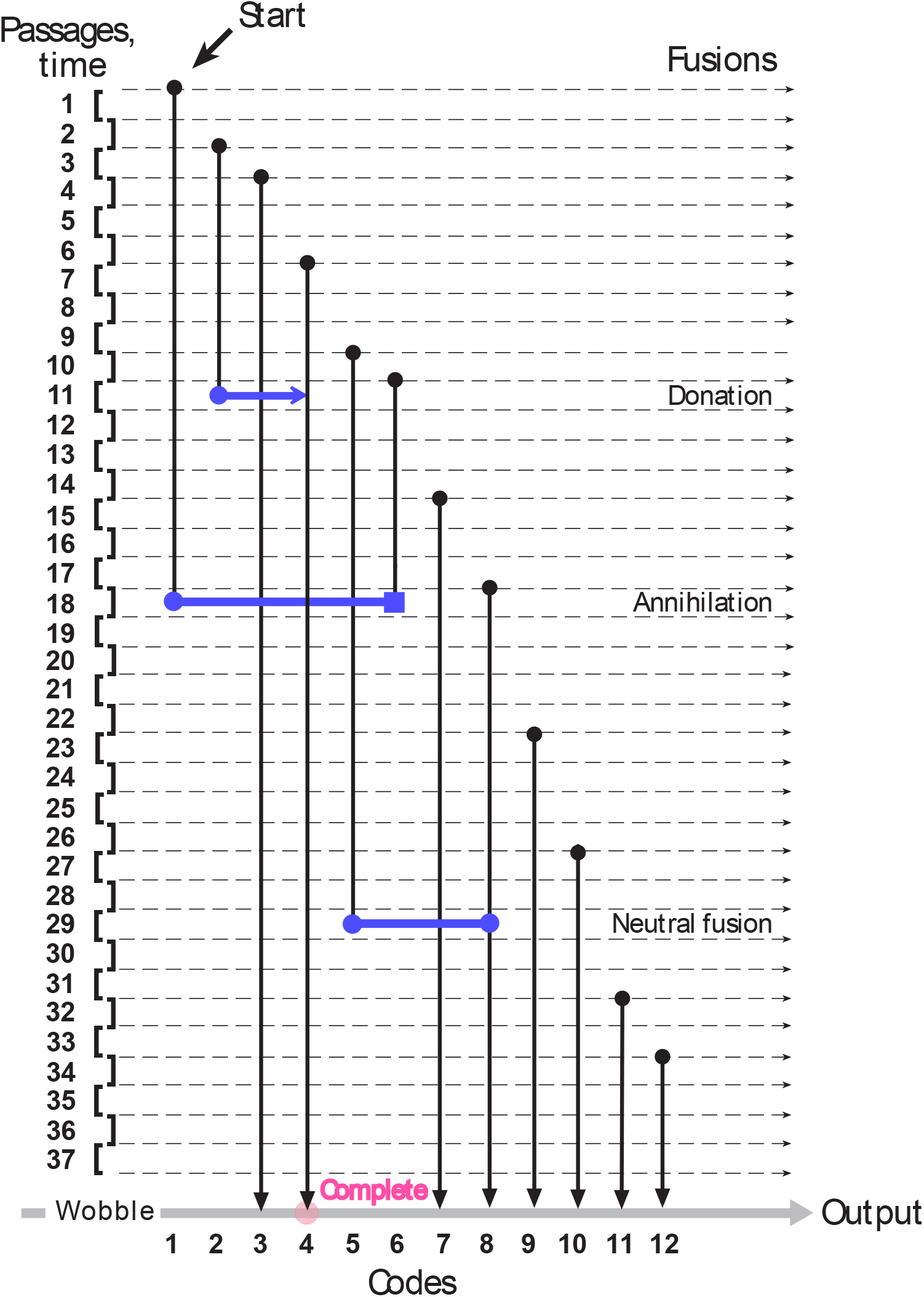
Code evolution in one simplified environment. For explanatory purposes, Fig. 7 shows only 37 passages (environments can have thousands) and 12 codes (environments can have hundreds). Passages are time for one event in coding evolution: individual passages vary stochastically, but are shown similarly for clarity. However, the mean passage is defined, marking time reasonably precisely. Each environment begins with a single table, labeled “Start”. New tables appear in Fig. 7 with constant probability per passage (Ptab), starting with a single arbitrary assignment (Pinit). At each passage, all current tables evolve by one step: a new assignment (Pinit), random (Prand) or SGC-like (1 – Prand), an assignment decay (Pdecay), or capture of an unassigned codon one mutation distant (Pmut). As soon as multiple tables arise, they can fuse with ({probability Pfus/passage}*{number of codes – 1}. Three kinds of fusions exist, each decreasing total codes. The vanishing code can contribute assignments to a recipient (“Donation”). The vanishing code can be lost, along with its incompatible recipient (“Annihilation”). The vanishing code can cause no change if all its assignments already exist in the recipient (“Neutral fusion”). An environment is completed (“Complete”) at a set time, or when the environment contains a code with desired properties (translucent red circle), such as encoding >= 20 assigned functions. Wobble (late Crick wobble, lCw) evolves later (Yarus 2021b), after fundamental assignments are made. The program usually reports (“Output”) averages of all codes, or alternatively, properties of best codes (e.g., most complete: translucent red circle). In Fig. 7, this best code (Code 4) gained new assignments by fusion.

In fact, reliance on all molecular mechanism is minimized here because, in part, ancient coding machineries are still uncertain. Thus c3-iCw avoids dependence on particular reactions - fusion advantages would likely be similar for any means of encoding in which fused parts of the coding table can develop separately.

### Summary of c3-lCw

Fusion joins partial codes which may have diverse encodings. Fusion must occur frequently if fusion is to alter code distributions (Fig. 1), because even a single coding table without such fusion may evolve to resemble the SGC (Yarus 2021b; Fig. 3C). Fusion profoundly alters coding environments (Fig. 2). Code fusions will likely be undirected, because fused codes cannot express a phenotype until after fusion itself. Thus, fusions that facilitate expansion toward the SGC will be accompanied by those that do not – incompatible coding annihilates both participants (Fig. 2, 7). A fusing population shrinks as fusion expands. Under these conditions, the major long-term fate of codes is entry into effective, innocuous or disastrous fusions – only 9.4% of ≈ 61000 codes that once existed survive in this late environment (750 passages, Fig. 2).

Such losses change the average survivor (Fig. 3, 4). A fusing population shows a crescendo of individuals (Fig. 3A) with near-complete codes (>= 20 functions) that also resemble majority encoding (0, 1, 2 or 3 differences from the SGC). This sustained rise is a specific product of fusion. It does not appear for similar coexisting multiple codes lacking fusion (Fig. 3B). In addition, SGC-like codes are everywhere fewer for a system in which no similar fusion partners exist (Fig. 3 C). Moreover, competence is delayed in both non-fusing systems (Fig. 3). Code fusions will be more readily selected: they present better codes sooner, more frequently, and in a smaller population, for a longer time.

Crescendo competence has a clear source: average misassignment is suppressed in fusing environments, compared to non-fusing ones (Fig. 4A). That is: variant codes have been selectively extinguished in unproductive fusion attempts (Fig. 2, 6). A fused population has made fewer erroneous random assignments (Fig. 4B), as well as fewer error-prone captures of mutational-related triplets (Fig. 4C). This requires only that selection favor codons with unique meanings. Thus fusion purification of a dominant encoding will be only slightly dependent on evolutionary details.

Environments evolving via code fusions produce competent codes more frequently (Yarus 2021b) than prior models. For example, ≈ 1% to ≈ 11% of the environments in Fig. 3A have <=3 misassignments with >= 20 functions - these frequencies seem within easy reach of selection for superior SGC translation. Or more specifically, exact (mis0 class) SGC codes exist in > 1 in 1000 environments (Prand = 0.05, Fig. 5), though only selection intrinsic to fusion has been applied. For this latter class, the only selection required is harvesting exact SGC’s among other codes. This exemplifies distribution fitness (Yarus 2021b), where a beneficial distributed property is selected from an excellent environmental minority. Nevertheless, self-refinement due to fusion still requires accurate assignments (Yarus 2021c), still plausibly estimated as <= 10% random (Fig. 5).

### C3-lCw is consistent with previous conjecture

Fusion was initially suggested in order to allow SGC incorporation of different forms of ordered assignments (Yarus 2021c); for example, related amino acids in the same coding table column or row. It was also suggested that code fusion could both speed appearance of the SGC and allow it to appear in a smaller population. These predictions are borne out in Fig. 2 and Fig. 3

### The surprising crescendo

Simple assumptions (fusions between incompatible partial codes are deleterious) yield striking convergence on an existing code, potentially selecting the SGC (Fig. 3, 4). Convergence suppresses error from varied sources (Fig. 4A, 4B, 4C), and thus is likely to be broadly applicable to other kinds of variation. But most significantly, complete and accurate codes accumulate continuously during a long era of prevalent fusion (Fig. 3A). The same events remove codes from the group of accurate codes as well as create them (Fig. 2), so particular accurate codes present at one time (Fig. 3A) differ from those earlier and later. Therefore, a population comprised of varying SGC-like codes persists, such codes increasing in the environment, until selection of a superior one among them. It is difficult to imagine a more effective protocol for selecting SGC-like encoding.

### Three definitions for two coding eras

In several important ways, SGC history can be divided into two eras, each with its own evolutionary rules (Knight et al. 2001).

### First definition: code expansion vs code stasis

The focus here is the early period of expansion of the code to its present scope, presumably selected via the ability of enlarged (more complete) codes to encode more competent peptides (Sengupta and Higgs 2015). The implied contrast is with a later period, after substantial completion of the SGC, when the code is approximately “frozen” (Crick 1968) because it must conserve a highly evolved prior proteome (Ardell and Sella 2002). However, even the later code evolves to some extent (Jukes and Osawa 1993), perhaps selectively changing late-evolved functions, like termination (Yarus 2021a).

### Second definition: RNA agents vs ribonucleopeptide agents

The first definition just above is approximately echoed in the distinction between an early era resting solely on the capabilities of RNA (Gilbert 1986) and a second later era of ribonucleopeptide agents (Fig. 6). This second transition must lie somewhere near Crick’s freezing point because aminoacyl-RNA synthetases themselves are very complex proteins (they have multiple specific substrate binding sites and perform multiple catalyses). A modern variety of such catalysts is only plausible when the code has become strongly constrained by extensive prior encoding.

### Third definition: primordial amino acids vs those from anabolism

Division into early and late eras is further reinforced by the prevalent belief that coding was initiated with readily available natural amino acids (Miller 1953), perhaps those most easily chemically synthesized under primitive conditions (Higgs and Pudritz 2009). The earliest amino acids are frequently specified as G, A, D and V (Ikehara 2009), all encoded by GNN codons (N is any nucleotide) in the final SGC (Higgs 2009). Later more complex metabolism permitted the addition of amino acids derived from evolved metabolites (Wong 1975; Taylor and Coates 1989; Di Giulio 2008). This third division is closely related to those above: while coding likely began on RNA (Yarus 2017), synthesis of the first encoded peptides provides new molecules with novel conformations and chemical groups for intermolecular interactions. Thus, in the period before freezing (first definition), early codes are rapidly expanding their repertoire of interactions by making peptides that then can enhance further code evolution. Progress toward codes so competent that they have hindered their own further evolution (first definition) is speeded by the appearance of structured riboucleoproteins (second definition; compare (Turk et al. 2011; Carter 2015; Müller et al. 2022)). Finally, a near-complete SGC arises in the era of the ribonucleopeptide translation apparatus (Fig. 6). This code participates in the biochemistry of the last common ancestor, including its varied assortment of specific aminoacyl-RNA synthetases, required to encode protein catalysts for a complex metabolism (Ribas de Pouplana 2020; Xavier et al. 2021).

### All two-era definitions may rely on the crescendo

Fig. 6 suggests that c3-lCw might delimit all eras. During the c3-lCw the code became so complex it might be called frozen; the c3-lCw is also when capable ribonucleopeptide catalysts became possible, and the c3-lCw also inaugurates the first epoch when an almost modern set of nucleopeptide monomers might exist.

Current evidence suggests that the most ancient organisms took the form of close laminations, with different organisms densely layered within 3.43 Gya stromatolites (Allwood et al. 2006), or close bundles of filaments 3.75 to 4.28 Gya in seafloor jasper (Papineau et al. 2022). Either formation permits close approach of cells of different origins and competency, and potentially encourages fusion of their codes.

Given these microscopic fossils, c3-lCw might have occurred ≈ 4 billion years ago, as proposed in Fig. 6.

### Late assignments to late amino acids

SGC coding triplets can be sequences extracted from ancient RNA binding sites for cognate amino acids (Yarus and Christian 1989; Rodin et al. 2011; reviewed in Yarus 2017). Such RNA-amino acid interactions presumably supplied assignments for the early coding tables that converge on the SGC in this work (Fig. 6). A counterargument sometimes offered (e.g., Koonin and Novozhilov 2017) is that metabolically complex, likely late-appearing amino acids like arginine (Janas et al. 2010) and tryptophan (Majerfeld et al. 2010) are among amino acids associated with triplet-containing RNA sites. But earlier, simpler amino acids like isoleucine also have prominent coding triplets (Lozupone et al. 2003). Moreover, assignment of triplets from RNA binding sites would probably continue in a later ribonucleopeptide era (Fig. 6). RNA sites for later, complex amino acids like arginine (Yarus and Christian 1989) would be used when advantageous. So, ribonucleopeptide enzymes could expand anabolism to complex amino acids, while their SGC assignments still utilized their fits to RNA binding sites.

### HGT and code universality

(Vetsigian et al. 2006) expresses Carl Woese’s conviction that early HGT (Horizontal Gene Transfer) was a crucial “innovation-sharing protocol”. Before the genome and inheritance, only creatures possessing similar genetic codes could share innovations evolved independently. Such communal innovation-sharing decreased variations in codes before accurate vertical inheritance had evolved. An independent learning model confirms that mutual genetic intelligibility via HGT, without vertical inheritance, could have universalized the SGT (Froese et al. 2018). Present work suggests that earlier, when only partial codes existed, HGT would speed the assembly of, purify the population of, and facilitate the selection of, SGC-like codes.

## Methods

The program that repeatedly evolves coding environments and reports code properties was written using the integrated development environment in Lazarus v. 2.20RC1 in its console mode, with the free Pascal FPC 3.2.2 compiler. Compiled mechanisms were run on a Dell XPS computer under 64-bit Microsoft Windows 10 @ 2.9 GHz on an Intel Core i9-8950HK CPU, using 32 GB of RAM.

The source used for all present calculations is Ctable20k.pas, available on request from the author. Results from the program, as tab-delimited files, were passed to Microsoft Excel 2016 for analysis and graphics. An example of spreadsheet analysis is also available on request.

Time is measured in cycles through code evolution, called passages. Passages, and other details of programed action, are depicted in Fig. 7. Assignment, decay and codon capture, which occur mutually exclusively during a passage through each single code, have been discussed previously (Yarus 2021b). Multiple codes and fusion are introduced in Fig. 1 to 5 above.

## Acknowledgements

Thanks to Norm Pace for an informative discussion of stromatolites, and to Bill McClain for helping clarify a draft ms.

